# GenVisR: Genomic Visualizations in R

**DOI:** 10.1101/043604

**Authors:** Zachary L. Skidmore, Alex H. Wagner, Robert Lesurf, Katie M. Campbell, Jason Kunisaki, Obi L. Griffith, Malachi Griffith

**Affiliations:** McDonnell Genome Institute, Washington University School of Medicine, St. Louis, MO, 63108, USA.; Department of Medicine, Washington University School of Medicine, St. Louis, MO, 63110, USA.; Siteman Cancer Center, Washington University School of Medicine, St. Louis, MO, 63110, USA.; Department of Genetics, Washington University School of Medicine, St. Louis, MO, 63110, USA.

## Abstract

**Summary:** Visualizing and summarizing data from genomic studies continues to be a challenge. Here we introduce the GenVisR package to addresses this challenge by providing highly customizable, publication-quality graphics focused on cohort level genome analyses. GenVisR provides a rapid and easy-to-use suite of genomic visualization tools, while maintaining a high degree of flexibility by leveraging the abilities of ggplot2 and bioconductor.

**Availability and Implementation:** GenVisR is an R package available via bioconductor (https://bioconductor.org/packages/GenVisR) under GPLv3. Support is available via GitHub (https://github.com/griffithlab/GenVisR/issues) and the Bioconductor support website.

## 1 Introduction

The continued development of massively parallel sequencing technologies has led to an exponential growth in the amount of genomic data produced (Kodama, et al., 2012). This growth has in turn enabled scientists to investigate increasingly large, cohort-level genomic datasets. Generating intuitive visualizations is a critical component in recognizing patterns and investigating underlying biological properties in cohorts under study. A significant bottleneck exists, however, between data generation and subsequent visualization and interpretation (Good, et al., 2014). Additionally, generating publication-quality figures for effective communication of these data typically requires *ad hoc* methods such as manual creation or extensive graphic manipulation with third party software. This process is both time intensive and difficult to automate/reproduce. Here we present GenVisR, a Bioconductor package to address these issues. GenVisR provides a user-friendly, flexible, and comprehensive suite of tools for visualizing complex genomic data in three categories (small variant, copy number alteration, and data quality).

## 2 Small Variant

The identification of small variants (SNVs and indels) within a genomic context is of paramount importance for the elucidation of the genetic basis of disease. Numerous tools and resources have been created to identify variants in sequencing data (Wang, et al., 2013). Conversely, few tools exist to visually display and summarize these variants across sample cohorts. Given a gene of interest, it is often useful to view variant occurrences in the context of the translated protein product (Zhang, et al., 2012). A variety of options exist to achieve this, however tools that offer both automation and flexibility to perform this task are lacking (Table S1) (Griffith, et al., 2015; Leiserson, et al., 2015; Nilsen, et al., 2012; Yin, et al., 2012; Zhou, et al., 2015). The function *lolliplot* was developed to allow for precise control over visualization options (**Figure 1A**). This includes the ability to choose Ensembl annotation databases for protein domain displays and to plot multiple tracks of mutations above and below the protein representation. Another common objective of genomic studies is to identify variant recurrence across multiple genes within a cohort. The GenVisR function *waterfall* was developed to calculate and rapidly illustrate the mutational burden of variants on both a gene and sample level, and further differentiates between variant types (**Figure 1B**). Visual detection of mutually exclusive genomic events at the variant level is emphasized in this visualization by arranging samples in a hierarchical fashion such that samples with mutations in the most recurrently mutated genes are ranked first. Finally, it is often informative to investigate the rate of transition and transversion mutations observed across a set of cases. For example, lung tumors originating from patients with a history of tobacco smoke exposure display a pattern of enrichment for C to A or G to T transversions (Govindan, et al., 2012). The function *TvTi* (transversion/transition) was developed to improve recognition of these types of patterns within a cohort.

**Fig. 1.**
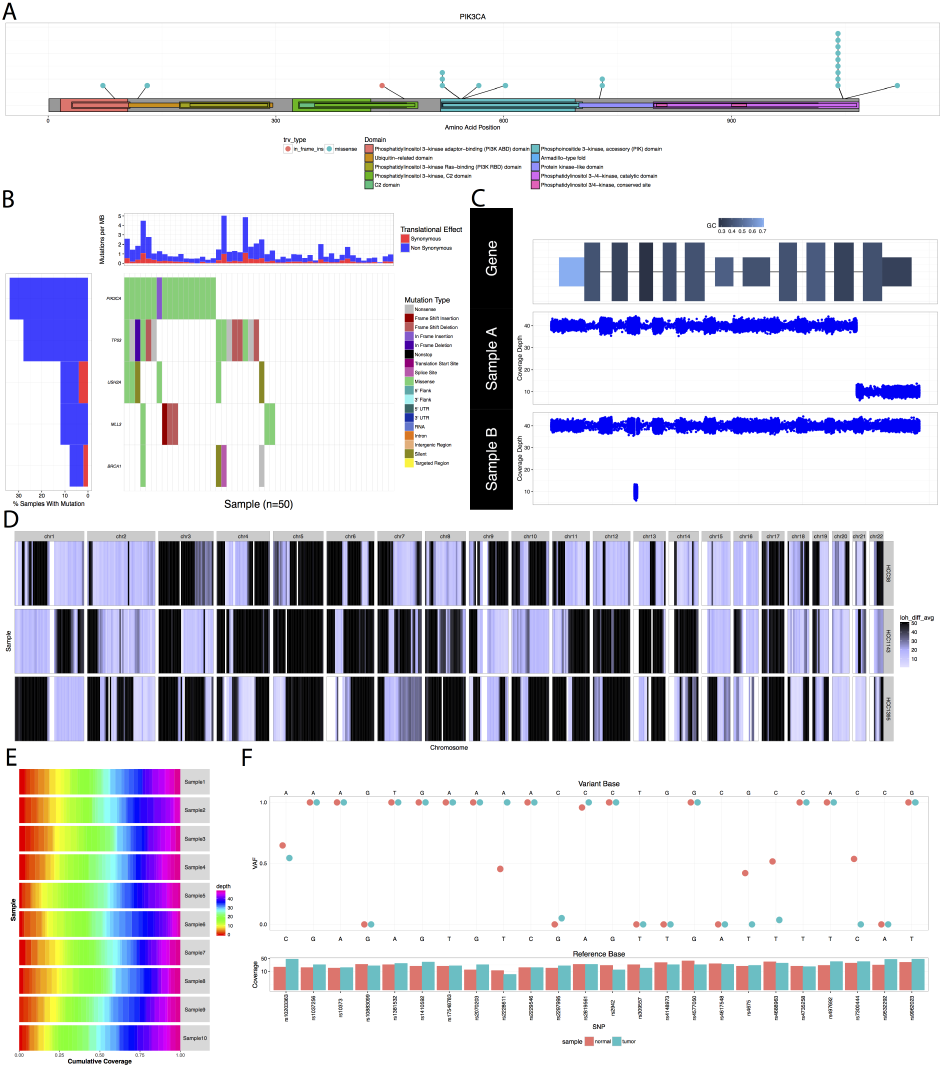
Selected representation of GenVisR visualizations A) Output from *lolliplot* for select TCGA breast cancer samples (Cancer Genome Atlas Network, 2012) shows two mutational hotspots in *PIK3CA* within the accessory and catalytic kinase domains. B) Output from *waterfall* shows mutations for five genes across 50 select TCGA breast cancer samples with mutation type indicated by colour in the grid and per sample/gene mutation rates indicated in the top and left sidebars. C) Output from *genCov* displays coverage (bottom plots) showing focal deletions in sample A (last exon) and B (second intron) within a gene of interest. GC content (top plot) is encoded via a range of colours for each exon. D) Output from *lohView* for HCC1395 (Griffith, et al., 2015), HCC38 and HCC1143 (Daemen, et al., 2013) breast cancer cell lines shows LOH events, across all chromosomes, shaded as dark blue. E) Output from *covBars* shows cumulative coverage for 10 samples indicating that for each sample, at least ~75% of targeted regions were covered at ≥ 35x depth. F) Output from *compIdent* for the HCC1395 breast cancer cell line (tumor and normal) shows variant coverage (bottom plot) and SNP allele fraction (main plot) indicating highly related samples. Note that 4/24 positions are discrepant and likely result from extensive LOH in this cell line.

## 3 Copy Number Alteration

Copy number alterations occurring within the genome are implicated in a variety of diseases (Beroukhim, et al., 2010). The function *GenCov* illustrates amplifications and deletions across one or more samples in a genomic region of interest (**Figure 1C**). A key feature of *GenCov* is the effective use of plot space, especially for large regions of interest, via the differential compression of various features (introns, exons, UTR) within the region of interest. For a broader view the function *cnView* plots copy number calls, and the corresponding ideogram, for an individual sample at the chromosome level. The function *cnSpec* displays amplifications and deletions on a still larger scale via copy number segments calls. This information is displayed as a heat map arranged in a grid indexed by chromosomes and samples. Alternatively, *cnFreq* displays the frequency of samples within a cohort that are observed to have copy number gains or losses at specific genomic loci. In addition to copy number changes, loss of heterozygosity (LOH) often plays an important role in genomic diseases. For example, in acute myeloid leukemia copy neutral LOH has been associated with shorter complete remission and worse overall survival (Gronseth, et al., 2015). The function *lohView* displays LOH regions observed within a cohort (**Figure 1D**) by plotting a sliding window mean difference in variant allele fractions for tumor and normal germline variants.

## 4 Data Quality

In genomic studies, the quality of sequencing data is of critical importance to the proper interpretation of observed variations.

Therefore, we provide a suite of functions focused on data quality assessment and visualization. The first of these, *covBars,* provides a framework for displaying the sequencing coverage achieved for targeted bases in a study (**Figure 1E**). A second function, *compIdent,* aids in the identification of mix-ups among samples that are thought to originate from the same individual (**Figure 1F**). This is achieved by displaying the variant allele fraction of SNPs in relation to each sample. By default, 24 biallelic “identity SNPs” (Pengelly, et al., 2013) are used to determine sample identity.

## 5 Example Usage

GenVisR was developed with the naïve R user in mind. Functions are well documented and have reasonable defaults set for optional parameters. To illustrate, creating **Figure 1B** was as simple as executing the *waterfall* function call on a standard MAF (version 2.4) file and choosing which genes to plot:

> genes = c(“PIK3CA”, “TP53”, “USH2”, “MLL3”, “BRCA1”) GenVisR::waterfall(x = maf_file, plotGenes=genes)

## 6 Conclusion

GenVisR provides features and functions for many popular genomic visualizations not otherwise available in a single convenient package (Table S1). By leveraging the abilities of ggplot2 (Wickham, 2009) it confers a level of customizability not previously possible. Virtually any aspect of a plot can be changed simply by adding an additional layer onto the graphical object. Thus, GenVisR allows for publication quality figures with a minimal amount of required input and data manipulation while maintaining a high degree of flexibility and customizability.

## Acknowledgements

The authors would like to thank Richard K. Wilson and Elaine R. Mardis for their encouragement and support in the creation of this work. We also thank Chris A. Miller and Dave Larson for their creative guidance.

## References

Beroukhim, R., et al. The landscape of somatic copy-number alteration across human cancers. Nature 2010;463(7283):899–905.

Cancer Genome Atlas Network. Comprehensive molecular portraits of human breast tumours. Nature 2012;490(7418):61–70.

Daemen, A., et al. Modeling precision treatment of breast cancer. Genome Biol 2013;14(10):R110.

Good, B.M., et al. Organizing knowledge to enable personalization of medicine in cancer. Genome Biol 2014;15(8):438.

Govindan, R., et al. Genomic landscape of non-small cell lung cancer in smokers and never-smokers. Cell 2012;150(6): 1121–1134.

Griffith, M., et al. Genome Modeling System: A Knowledge Management Platform for Genomics. PLoS Comput Biol 2015;11(7):e1004274.

Gronseth, C.M., et al. Prognostic significance of acquired copy-neutral loss of heterozygosity in acute myeloid leukemia. Cancer 2015;121(17):2900–2908.

Kodama, Y., et al. The Sequence Read Archive: explosive growth of sequencing data. Nucleic Acids Res 2012;40(Database issue):D54–56.

Leiserson, M.D., et al. MAGI: visualization and collaborative annotation of genomic aberrations. Nat Methods 2015;12(6):483–484.

Nilsen, G., et al. Copynumber: Efficient algorithms for single‐ and multi-track copy number segmentation. BMC Genomics 2012;13:591.

Pengelly, R.J., et al. A SNP profiling panel for sample tracking in whole-exome sequencing studies. Genome Med 2013;5(9):89.

Wang, Q., et al. Detecting somatic point mutations in cancer genome sequencing data: a comparison of mutation callers. Genome Med 2013;5(10):91.

Wickham, H. ggplot2: Elegant Graphics for Data Analysis. Springer Publishing Company, Incorporated; 2009.

Yin, T., Cook, D. and Lawrence, M. ggbio: an R package for extending the grammar of graphics for genomic data. Genome Biol 2012;13(8):R77.

Zhang, J., et al. The genetic basis of early T-cell precursor acute lymphoblastic leukaemia. Nature 2012;481(7380):157–163.

Zhou, X., et al. Exploring genomic alteration in pediatric cancer using ProteinPaint. Nat Genet 2015;48(1):4–6.

